# CHD8 Regulates Cellular Homeostasis and Neuronal Function Genes Across Multiple Models of *CHD8* Haploinsufficiency

**DOI:** 10.1101/375238

**Authors:** A. Ayanna Wade, Kenneth Lim, Rinaldo Catta-Preta, Alex S. Nord

## Abstract

The packaging of DNA into chromatin determines the transcriptional potential of cells and is central to eukaryotic gene regulation. Recent sequencing of patient mutations has linked *de novo* loss-of-function mutations to chromatin remodeling factors with specific, causal roles in neurodevelopmental disorders. Characterizing cellular and molecular phenotypes arising from haploinsufficiency of chromatin remodeling factors could reveal convergent mechanisms of pathology. Chromodomain helicase DNA binding protein 8 (*CHD8*) encodes a chromatin remodeling factor gene and has among the highest *de novo* loss-of-function mutations rates in patients with autism spectrum disorder (ASD). Mutations to *CHD8* are expected to drive neurodevelopmental pathology through global disruptions to gene expression and chromatin state, however, mechanisms associated with CHD8 function have yet to be fully elucidated. We analyzed published transcriptomic and epigenomic data across *CHD8 in vitro* and *in vivo* knockdown and knockout models to identify convergent mechanisms of gene regulation by CHD8. We found reproducible high-affinity interactions of CHD8 near promoters of genes necessary for basic cell functions and gene regulation, especially chromatin organization and RNA processing genes. Overlap between CHD8 interaction and differential expression suggests that reduced dosage of *CHD8* directly relates to decreased expression of these genes. In addition, genes important for neuronal development and function showed consistent dysregulation, though there was a reduced rate and decreased affinity for CHD8 interactions near these genes. This meta-analysis verifies CHD8 as a critical regulator of gene expression and reveals a consistent set of high affinity CHD8 interaction targets observed across human and mouse *in vivo* and *in vitro* studies. Our findings highlight novel core functions of CHD8 and indicate direct and downstream gene regulatory impacts that are likely to be associated with neuropathology underlying CHD8-associated neurodevelopmental disorder.

## INTRODUCTION

Recent genetic studies suggest that single copy loss-of-function mutations to chromatin remodeling genes significantly contribute to autism spectrum disorder (ASD) neurobiology, presumably through disruptions to transcriptional regulation in the developing and mature brain (De Rubeis et al. 2014, Iossifov et al. 2014, Parikshak et al. 2013, O’Roak et al. 2012a, O’Roak et al. 2012b, Sanders et al. 2015, Vissers et al. 2016). The gene encoding chromodomain helicase DNA binding protein 8 (*CHD8*) has one of the highest observed mutation rates in sporadic ASD (O’Roak et al. 2012a, Barnard et al. 2015, Krumm et al. 2014), and mutations to *CHD8* have also been identified in cases from schizophrenia and intellectual disability cohorts (Tatton-Brown et al. 2017, McCarthy et al. 2014). Patients that carry *CHD8* mutations typically meet stringent qualifications for ASD diagnosis, and frequently present with comorbid features including macrocephaly, cognitive impairment, distinct craniofacial morphology, and gastrointestinal disturbances (Bernier et al. 2014). Knockdown or haploinsufficiency of *Chd8* in animal models has recapitulated specific neuroanatomical, gastrointestinal, cognitive, and behavioral phenotypes observed in patients (Sugathan et al. 2014, Gompers et al. 2017, Katayama et al. 2016, Platt et al. 2017), though reported phenotypes vary across models. Considering the relevance of *CHD8* mutations in neurodevelopmental disorders and the positive findings of relevant phenotypes in model systems, characterizing the convergent patterns of CHD8 genomic interactions and transcriptional outcomes caused by *CHD8* haploinsufficiency across studies could significantly advance understanding of core pathophysiology in patients carrying *CHD8* mutations and, potentially, reveal generalized chromatin-associated cellular mechanisms underlying neurodevelopmental disorders.

CHD8 belongs to a family of ATP-dependent chromatin remodelers (Hall and Georgel 2007, Marfella and Imbalzano 2007, Hargreaves and Crabtree 2011). CHD family proteins are distinguished by tandem chromodomains predicted to enable these proteins to bind histones (Flanagan et al. 2005). As some CHD proteins demonstrate chromatin remodeling activity (Hall and Georgel 2007, McKnight et al. 2011, Tong et al. 1998), CHD8 has been speculated to drive ASD-associated changes in neurodevelopmental gene expression by targeting and remodeling chromatin at specific promoters and enhancers (Sugathan et al. 2014, Cotney et al. 2015, Ceballos-Chavez et al. 2015). This is supported by evidence that CHD8 can reposition nucleosomes *in vitro* and in mammalian cell culture (Thompson et al. 2008), and that loss of *Chd8* in *in vitro* and *in vivo* models dysregulates ASD-associated and Chd8-target gene expression (Sugathan et al. 2014, Gompers et al. 2017, Katayama et al. 2016, Cotney et al. 2015). Several mechanisms have been suggested to underlie CHD8 binding specificity, including targeting through histone modifications associated with open chromatin (Sugathan et al. 2014, Cotney et al. 2015, Yuan et al. 2007, Rodriguez-Paredes et al. 2009) and recruitment through protein-protein interactions (Thompson et al. 2008, Yuan et al. 2007, Rodriguez-Paredes et al. 2009, Ishihara et al. 2006, Nishiyama et al. 2009, Shen et al. 2015, Fang et al. 2016). However, the mechanisms by which CHD8 directly regulates target gene expression, whether CHD8 targets cell- and stage-specific genes in the developing brain, and which patterns of transcriptional dysregulation are due to direct effects versus downstream or secondary changes to CHD8 regulation remain unresolved.

There are a growing number of studies that have explored the role of CHD8 in neurodevelopment, providing the opportunity to test for core features of CHD8 genomic interactions and transcriptomic dysregulation associated with *CHD8* haploinsufficiency. Published studies have encompassed both *in vitro* and *in vivo* systems with shRNA knockdown or genetic mutation of *CHD8.* Despite the variety of models there appear to be general patterns of neurodevelopmental disruption caused by reduced *CHD8* expression, characterized by impacts to cellular proliferation, neuronal differentiation, and synaptic function. However, discrepancies between cellular and behavioral findings make it difficult to reconcile core features of cellular pathology. While published models of *CHD8* haploinsufficiency vary considerably, nearly all such studies have leveraged genomic approaches to determine the impact of *CHD8* haploinsufficiency on gene expression. Many have also examined CHD8 interaction targets genome-wide. The methods used for these experiments, RNA sequencing (RNA-seq) and chromatin immunoprecipitation followed by sequencing (ChIP-seq) can generate comparable, unbiased, and quantitative data enabling direct comparisons of results across models and studies. We hypothesized that meta-analysis of these datasets using the same computational methods may capture the consistent patterns of transcriptional pathology associated with *CHD8* haploinsufficiency and reveal constitutive and model-specific genomic interaction patterns of CHD8.

Here, we re-analyzed published RNA- and ChIP-seq data from *CHD8 in vitro* and *in vivo* mouse and human models, and built an online user interface to enable customizable data analysis and visualization across transcriptomic studies of *CHD8* haploinsufficiency. Across studies, we found a reproducible set of high-affinity CHD8 interaction target genes important for cellular homeostasis, with many datasets additionally exhibiting downregulated gene expression of these targets. We also found a secondary signature of transcriptional dysregulation of genes important for neuronal development and function consistent with *CHD8* haploinsufficiency. The findings of this meta-analysis indicate evolutionarily-conserved functions of CHD8, with reductions in *CHD8* expression directly and indirectly altering transcription of genes critical for cellular homeostasis and neurodevelopment in mouse and human models.

## MATERIALS AND METHODS

### *CHD8* genomic datasets

Next generation sequencing datasets generated from *CHD8* studies were identified through a literature search of publications featuring the keyword “CHD8” in the PubMed and Gene Expression Omnibus (GEO) databases. Raw data from publications that featured RNA-seq or ChIP-seq analysis were downloaded from GEO with the exception of three publications that hosted raw data on DDBJ (Katayama et al. 2016) and SRA (Platt et al. 2017, Wilkinson et al. 2015). A total of twelve publications corresponding to 289 sequencing libraries were included in the analysis. Libraries from Cotney et al. (2015) generated from fetal brain and libraries from Han et al. (2017) designed for analysis of alternative splicing were not included in the analysis. All data included were stated to be compliance with respective animal care and use committees at time of original publication.

### RNA-seq analysis

RNA-seq computational analysis was performed following an established pipeline using standard software, as described previously (Gompers et al. 2017). Briefly, unaligned sequencing reads were assessed for general quality using FastQC (Version 0.11.2) and aligned to the mouse (mm9) or human (GRCh37) reference genome using STAR (Version 2.5.2b, Dobin et al. 2013). Aligned reads mapping to genes according to the mm9 genes.gtf or to gencode.v19.annotation.gtf were counted at the gene level using subreads featureCounts (Version 1.5.0-p1, Liao et al. 2014). Overall data quality, including testing for GC-bias, gene body coverage bias, and proportion of reads in exons was further assessed using RSeQC (Version 2.6.4, Wang et al. 2012). Raw gene count data and sample information as reported in the respective repositories were used for differential expression analysis using edgeR (Version 3.4.4, Robinson et al. 2010). Genes with at least 1 count per million were included in a general linearized model using a sequencing-run factor-based covariate with genotype or knockdown as the variables for testing. For some datasets additional covariates were included if described in the original publication. Where possible, overall patterns of differentially expressed genes were compared to the original publication to ensure consistency in results. Normalized expression levels were generated using the edgeR rpkm function. Normalized log2(RPKM) values were used for plotting summary heatmaps and for expression data of individual genes. Variation in sequencing depth and intra-study sample variability partially account for differences in sensitivity and power across studies and likely drive some of the differences observed across studies, including the total number of differentially expressed genes. To capture an inclusive set of differentially expressed genes (DEGs), DEGs were defined by uncorrected p-values < 0.05. DEG sets were used for gene set enrichment analysis for Gene Ontology terms.

### ChIP-seq analysis

ChIP-seq analysis was also performed using an established pipeline and standard methods, as reported before (Gompers et al. 2017). Briefly, unaligned sequencing reads were assessed for general quality using FastQC and mapped to the mouse (mm9) or human (hg19) genome using BWA (Version 0.7.13, Li and Durbin 2009). Significant peaks with a p-value of < 0.0001 were identified using MACS2 (Version 2.1.0, Feng et al. 2011) with model-based peak identification and local significance testing disabled. Test datasets were analyzed comparing each individual ChIP-seq experiment to matched input or IgG controls. Input and IgG libraries were analyzed using the same approach to test for technical artifacts that could confound ChIP-seq results generally following a previously reported quality control strategy (Marinov et al. 2014). Enriched regions from IP and control datasets were annotated to genomic features using custom R scripts and the combined UCSC and RefSeq transcript sets for the mouse or human genome build. CHD8 target genes were assigned by peak annotation to transcript start site (TSS) or to the nearest TSS for distal peaks. HOMER was used to perform *de novo* motif discovery with default parameters (Version 4.7, Heinz et al. 2010). Where possible, we verified that results from ChIP-seq reanalysis were consistent with original publication.

### Gene ontology enrichment

The goseq R package (Version 1.30.0, Young et al. 2010) was used to test for enrichment of gene ontology terms while correcting for gene length. Analysis included GO Biological Process, Molecular Function, and Cellular Component annotations and required a minimal node size, or number of genes annotated to GO terms, of 20. The internal ‘weight01’ testing framework and Fishers test was used to account for multiple testing comparisons. Down- and upregulated genes were examined separately for RNA-seq GO analysis using a goseq FDR < 0.05 cutoff. ChlP-seq gene sets for the GO analysis were analyzed for experimental and control libraries separately also with a goseq FDR < 0.05 cutoff. Test gene sets for DEGs and CHD8 interaction targets were compared against a background set of expressed genes based on the minimum read-count cutoffs for each dataset for DEGs or a background set of all conserved mouse-human genes identified across RNA-seq datasets for CHD8 target genes. Heatmaps showing positive log2(expected/observed) values were plotted for GO terms for data visualization.

### Code and data availability and additional analysis visualization

Data that support the findings of this study are available from the corresponding author upon request. Accession numbers in parentheses and DOIs for all published gene sets used in enrichment analysis:

Ceballos-Chavez et al. (GSE62428): https://dx.doi.org/10.1371/journal.pgen.1005174;

Cotney et al. (GSE57369): https://dx.doi.org/10.1038/ncomms7404;

de Dieuleveult et al. (GSE64825): https://dx.doi.org/10.1038/nature16505;

Durak et al. (GSE72442): https://dx.doi.org/10.1038/nn.4400;

Gompers et al. (GSE99331): https://dx.doi.org/10.1038/nn.4592;

Katayama et al. (DRA003116): https://dx.doi.org/10.1038/nature19357;

Platt et al. (PRJNA379430): https://dx.doi.org/10.1016/j.celrep.2017.03.052;

Shen et al. (GSE71183, GSE71185): https://dx.doi.org/10.1016/j.molcel.2015.10.033;

Sugathan et al. (GSE61492): https://dx.doi.org/10.1073/pnas.1405266111;

Wang et al. 2015 (GSE71594): https://dx.doi.org/10.1186/s13229-015-0048-6;

Wang et al. 2017 (GSE85417): https://dx.doi.org/10.1186/s13229-017-0124-1;

Wilkinson et al. (PRJNA305612): https://dx.doi.org/10.1038/tp.2015.62;

Expanded results of the meta-analysis reported here are available from the interactive web server at https://nordlab.shinyapps.io/rnabrowser/. ChlP-seq datasets are available as UCSC TrackHubs for upload to the UCSC Genome Browser. All custom scripts for data processing and analysis are available at https://github.com/NordNeurogenomicsLab/.

## RESULTS

### Consistent patterns of transcriptional pathology associated with *CHD8* haploinsufficiency

We reanalyzed a total of 240 RNA sequencing libraries corresponding to 10 studies of *CHD8* knockdown or heterozygous mutation (**Table 1**). Almost all datasets represented neuronal model systems except for one dataset using an acute myeloid leukemia cell line (Shen et al. 2015). Analysis of all datasets was performed using the same pipeline with quality control steps and study-specific exceptions for consistency and covariate and batch structure as described in original publication (**Figure 1A**). Unsurprisingly, relative gene expression levels varied widely across studies, with principle components of variation dominated by species of origin and experiment (**Figure 1B**). However, pairwise comparisons between DEGs from individual datasets revealed specific similarities in gene expression changes. For example, comparison of DEGs at the p < 0.01 cutoff level between the Gompers et al. (2017) and the Sugathan et al. (2014) datasets revealed a strong positive correlation in direction of differential gene expression, where genes that were significantly up- or down-regulated in one dataset followed the same pattern in the other (**Figure 1C**). Further pairwise comparisons between studies and expression for specific genes can be done using our interactive web browser available at https://github.com/NordNeurogenomicsLab/. This interactive resource allows for analysis of principle components, differential expression of individual genes, and overall differential expression patterns for all included datasets (**Supplemental Figure 1**). New data from *CHD8* models will be added to this site as they are published and available.

**Figure 1.**
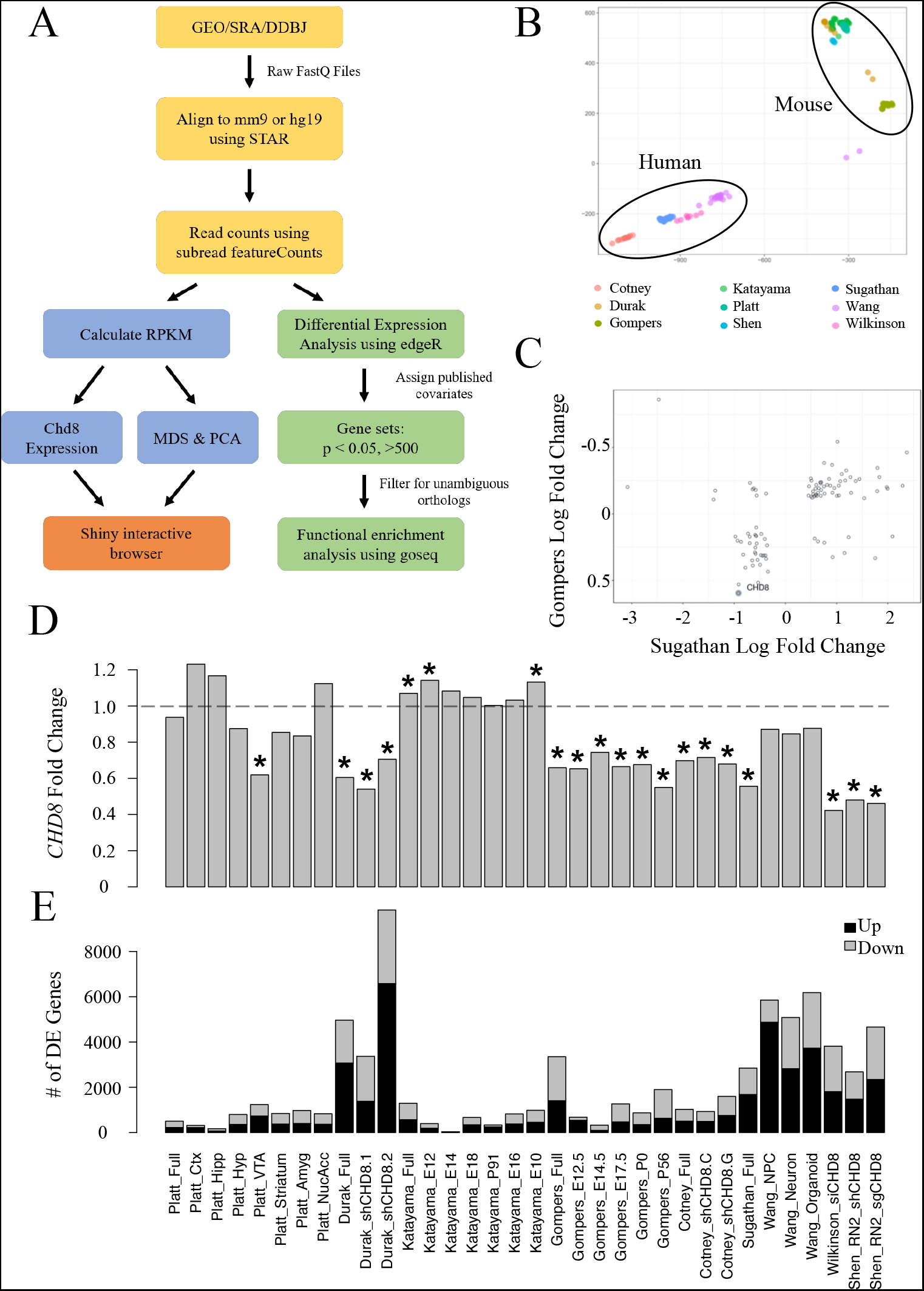
Differential gene expression across *CHD8* models. (A) RNA-seq data analysis pipeline. (B) PCA showing similarity in gene expression according to species. (C) Correlation between the Gompers et al. 2017 and Sugathan et al. 2014 RNA-seq datasets (p < 0.01) generated from the Shiny web browser. (D) Change in *CHD8* mRNA across models (p < 0.05). (E) Differential expression genes count across models (p < 0.05).

**Table 1.**
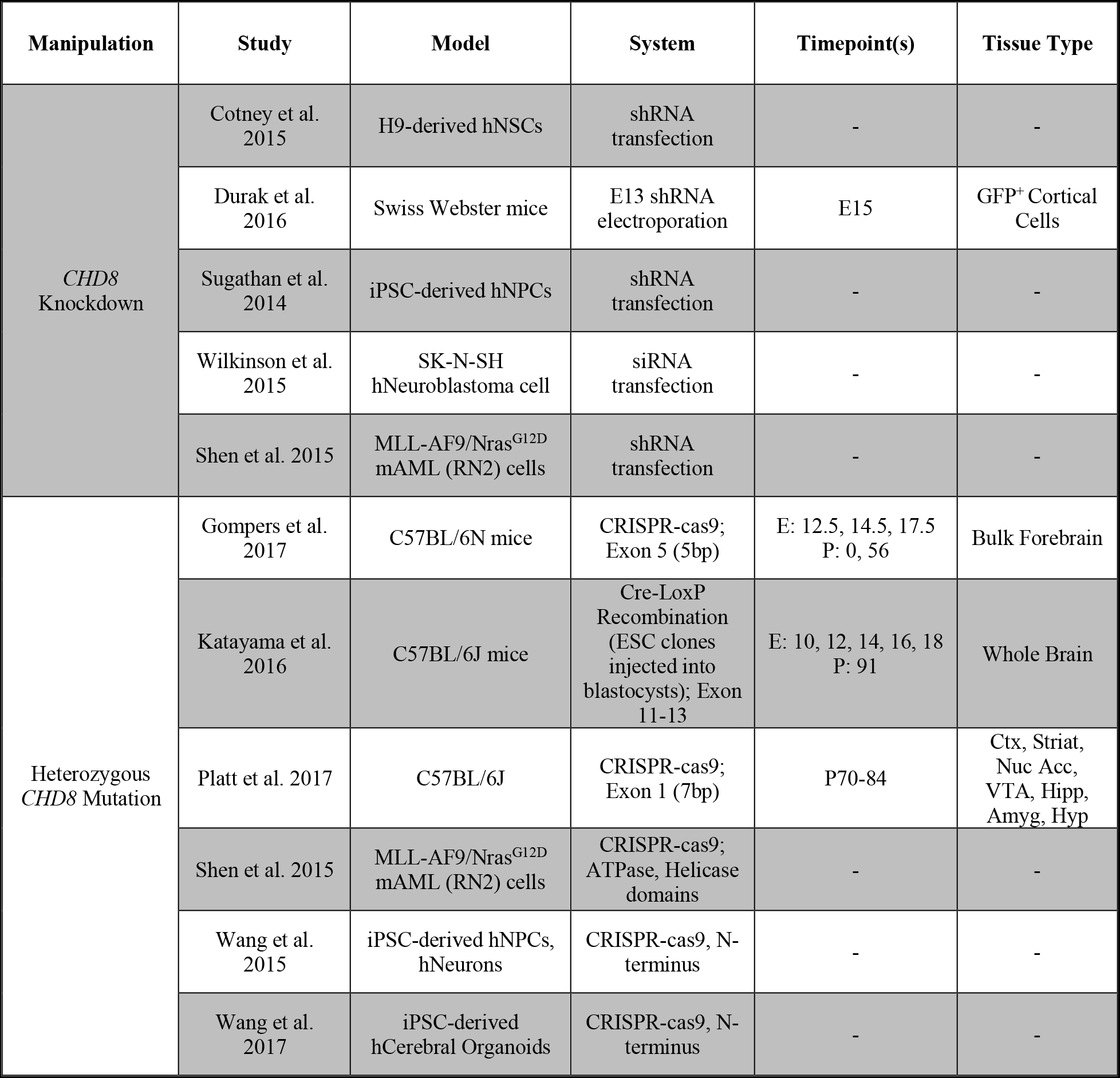
Summary of RNA-seq datasets included in the *CHD8* model reanalysis; h - human, NSCs - neural stem cells, iPSC - induced pluripotent stem cell, NPC - neural progenitor cell, E - embryonic day, P - postnatal day, Ctx - Prefrontal Cortex, Striat - Dorsal Striatum, Nuc Acc - Nucleus Accumbens, VTA - Ventral Tegmental Area, Hipp - Hippocampal Formation, Amyg - Amygdala, Hyp - Lateral Hypothalamus.

| Manipulation | Study | Model | System | Timepoint(s) | Tissue Type |
| --- | --- | --- | --- | --- | --- |
| *CHD8* Knockdown | Cotney et al. 2015 | H9-derived hNSCs | shRNA transfection | - | - |
|  | Durak et al. 2016 | Swiss Webster mice | E13 shRNA electroporation | E15 | GFP<sup>+</sup> Cortical Cells |
|  | Sugathan et al. 2014 | iPSC-derived hNPCs | shRNA transfection | - | - |
|  | Wilkinson et al. 2015 | SK-N-SH hNeuroblastoma cell | siRNA transfection | - | - |
|  | Shen et al. 2015 | MLL-AF9/Nras<sup>G12D</sup> mAML (RN2) cells | shRNA transfection | - | - |
| Heterozygous *CHD8* Mutation | Gompers et al. 2017 | C57BL/6N mice | CRISPR-cas9; Exon 5 (5bp) | E: 12.5, 14.5, 17.5<br>P: 0, 56 | Bulk Forebrain |
|  | Katayama et al. 2016 | C57BL/6J mice | Cre-LoxP Recombination (ESC clones injected into blastocysts); Exon 11-13 | E: 10, 12, 14, 16, 18<br>P: 91 | Whole Brain |
|  | Platt et al. 2017 | C57BL/6J | CRISPR-cas9; Exon 1 (7bp) | P70-84 | Ctx, Striat, Nuc Acc, VTA, Hipp, Amyg, Hyp |
|  | Shen et al. 2015 | MLL-AF9/Nras<sup>G12D</sup> mAML (RN2) cells | CRISPR-cas9; ATPase, Helicase domains | - | - |
|  | Wang et al. 2015 | iPSC-derived hNPCs, hNeurons | CRISPR-cas9, N-terminus | - | - |
|  | Wang et al. 2017 | iPSC-derived hCerebral Organoids | CRISPR-cas9, N-terminus | - | - |

Considering expression of *CHD8* itself, most knockdown and heterozygous knockout models resulted in a 50-60% significant decrease in mRNA (**Figure 1D**). However, published data from some models only showed a subtle decrease or even a significant increase in *CHD8.* We verified that these findings were consistent with originally published RNA-seq data. The absence of reduced *CHD8* mRNA expression for some studies raises questions regarding what expectations should be for gene dosage models. We note that protein level validation of *CHD8* dosage decrease was performed in all original publications to confirm *CHD8* haploinsufficiency in each model but considering the use of a number of different and unvalidated CHD8 antibodies across the studies, it is impossible to compare the protein validation results.

As expected, across all studies there were upregulated and downregulated genes passing stringent thresholds, though the numbers of DEGs varied widely. Consistent with the original publications, this re-analysis demonstrates CHD8 has direct or indirect roles in both facilitating and repressing gene expression (**Figure 1E**). Large differences in number and effect size of differentially expressed genes across studies may be a result of differences in experimental design, impact of knockdown and knockout on *CHD8* dosage, methods, and statistical sensitivity related to intra-study sample variability and sequencing depth. Variability in gene expression could also be due to differences in sensitivity to *CHD8* dosage between developmental stages and type of model used to carry out these experiments.

### Changes in *CHD8* expression affect genes important for cellular homeostasis and neuronal function

To examine patterns of transcriptional dysregulation associated with knockdown or heterozygous mutation to *CHD8*, we performed gene set enrichment analysis of Gene Ontology (GO) terms including datasets with at least 500 differentially expressed genes (**Figure 2**). Specific GO terms were chosen based on observed changes or interest to the field based on previous findings (for example, “canonical Wnt signaling pathway”). The full list of GO terms and relative enrichment is also provided (**Supplementary Figure 2**). While relatively small numbers of individual genes showed overlapping significant changes in expression across pairwise study comparisons, we found strong correlation in DEG functional groups across studies at the gene set level. This analysis identified two general signatures of differential gene expression across published models. The first signature was characterized by downregulation of genes annotated to terms related to the regulation of chromatin, transcription, and RNA processing, which we refer to as general cellular homeostasis (homeostasis) genes. As a whole, these are genes that do not exhibit cell specificity and are necessary for basic cell functions, such as chromatin organization, transcription and translation, and mitosis. This includes terms such as “RNA splicing,” “regulation of gene expression,” and “cell cycle.” The second signature encompassed terms related to neuronal development, maturation, and function, including terms associated with neural progenitor activity and lineage specification, synaptic function, and cell adhesion. We refer to these genes as neuronal function (neuronal) genes, and these genes showed both down- and up-regulation depending on the model. Examples of these terms include “neuron differentiation,” “axon guidance,” and “cell adhesion.”

**Figure 2.**
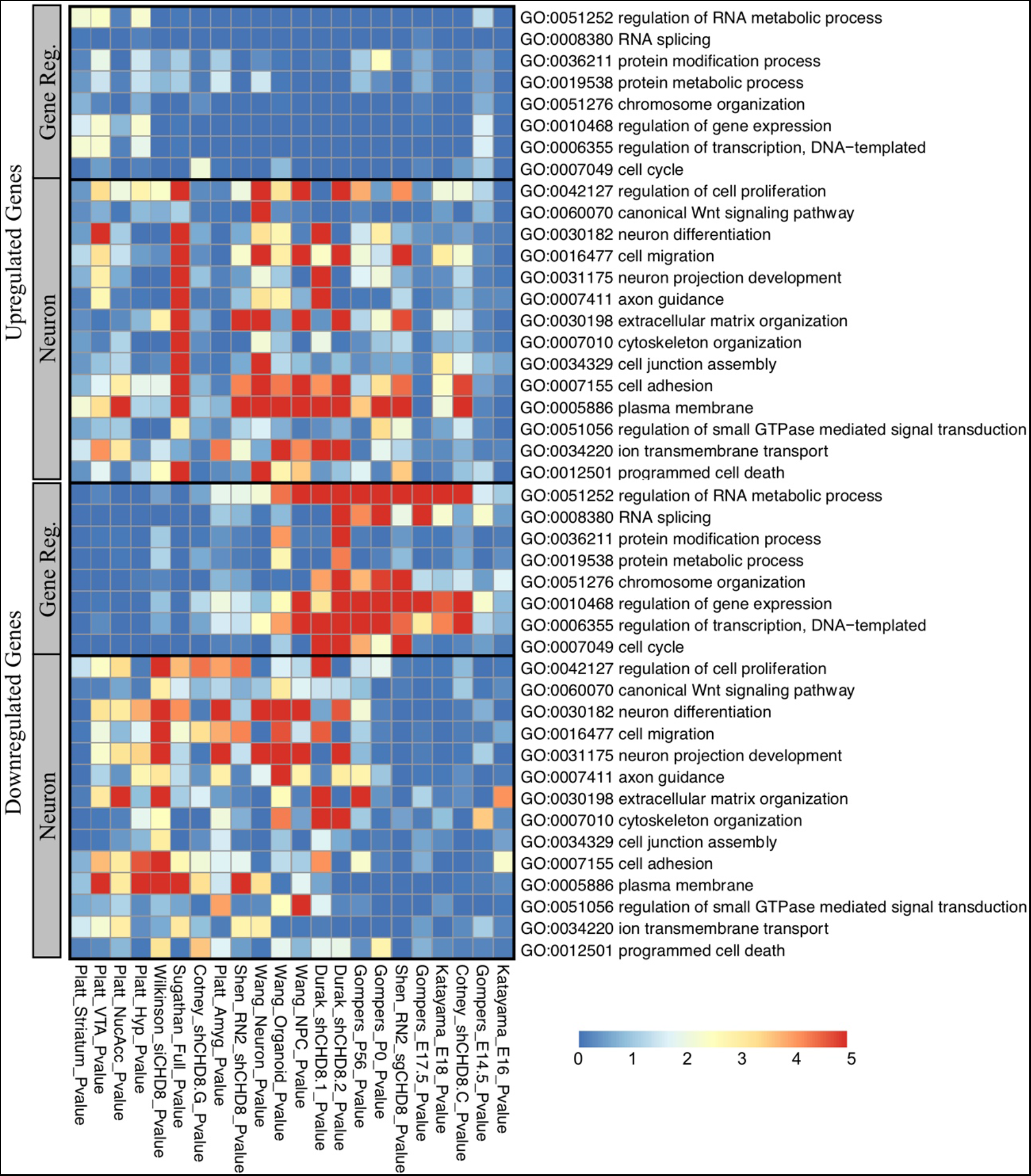
Enrichment of gene regulation and neurodevelopmental ontology terms in Up- and Down-regulated datasets having greater than 500 differentially expressed genes at the p < 0.05 level. (Top) Upregulated gene ontology enrichment. (Bottom) Downregulated gene ontology enrichment.

GO analysis showed different patterns of expression between upregulated and downregulated genes. Upregulated genes were mainly enriched for neuronal terms across models (**Figure 2, top**). In contrast, downregulated genes were enriched in homeostatic and neuronal terms in distinct patterns (**Figure 2, bottom**). Of 22 datasets, around 7 had neuronal terms enriched, 7 had homeostatic terms enriched, and 8 had a combination of both. The trend of enrichment of these signatures showed some correlation to the model system used in each study. *In vitro* models were more likely to have neuronal terms represented while in *vivo* models were more likely to have both, or only homeostatic terms, represented. There is also some indication that *in vivo* models of postnatal brain were more likely to have enrichment of neuronal terms while models of embryonic brain were more likely to have enrichment of homeostasis terms, but this remains a preliminary assessment requiring more robust data across developmental stages. Hierarchical clustering of all RNA-seq datasets reinforced this pattern, though enrichment of these signatures was weaker for datasets with fewer than 500 differentially expressed genes (**Supplementary Figure 3**). We note that there were also GO terms enriched only in individual datasets (**Supplementary Figure 2**). Overall, our results suggest that *CHD8* knockdown or heterozygous knockout consistently influences homeostatic and neuronal pathways, which are likely to drive the cellular, anatomical, and behavioral pathology reported in studies of *CHD8* haploinsufficiency.

CHD8-DNA interactions occur throughout the genome enriched for promoters

We reanalyzed a total of 49 ChIP-seq sequencing libraries from 8 studies of CHD8 genomic interaction patterns (**Table 2**). Analyzed datasets represented both neuronal and nonneuronal model systems. We included both *in vivo* tissue preparations and *in vitro* culture models from neuronal and non-neuronal fate cells to allow additional examination of tissue or cell-type specificity of CHD8 interactions. Half of the datasets were generated from bulk mouse tissue at adult (3 studies; Gompers et al. 2017, Katayama et al. 2016, Platt et al. 2017) and embryonic (2 studies; Katayama et al. 2016, Cotney et al. 2015) timepoints allowing for investigation of CHD8 interactions *in vivo* across time. The remaining data were generated from cellular models, with two studies using human neuronal lineage cells (Sugathan et al. 2014, Cotney et al. 2015), two using mouse or human cancer cell lines (Ceballos-Chavez et al. 2015, Shen et al. 2015), and one using mouse embryonic stem cells (De Dieuleveult et al. 2016). ChIP-seq data were analyzed using the same steps for immunoprecipitated, or experimental, and control data in our analysis pipeline (**Figure 3A**). There was large variation in number of called peaks, likely due to experimental design and technical differences (**Figure 3B**). Eleven of the control ChIP-seq libraries were found to have more than 250 called peaks with strong promoter enrichment (**Figure 3B-C**), suggesting some level of technical artifact associated with chromatin preparation (Marinov et al. 2014). Considering these experimental issues, control ChIP-seq libraries with >250 peaks were included in the analysis to test for similarity between technical artifacts and CHD8 immunoprecipitated signatures in these datasets.

**Figure 3.**
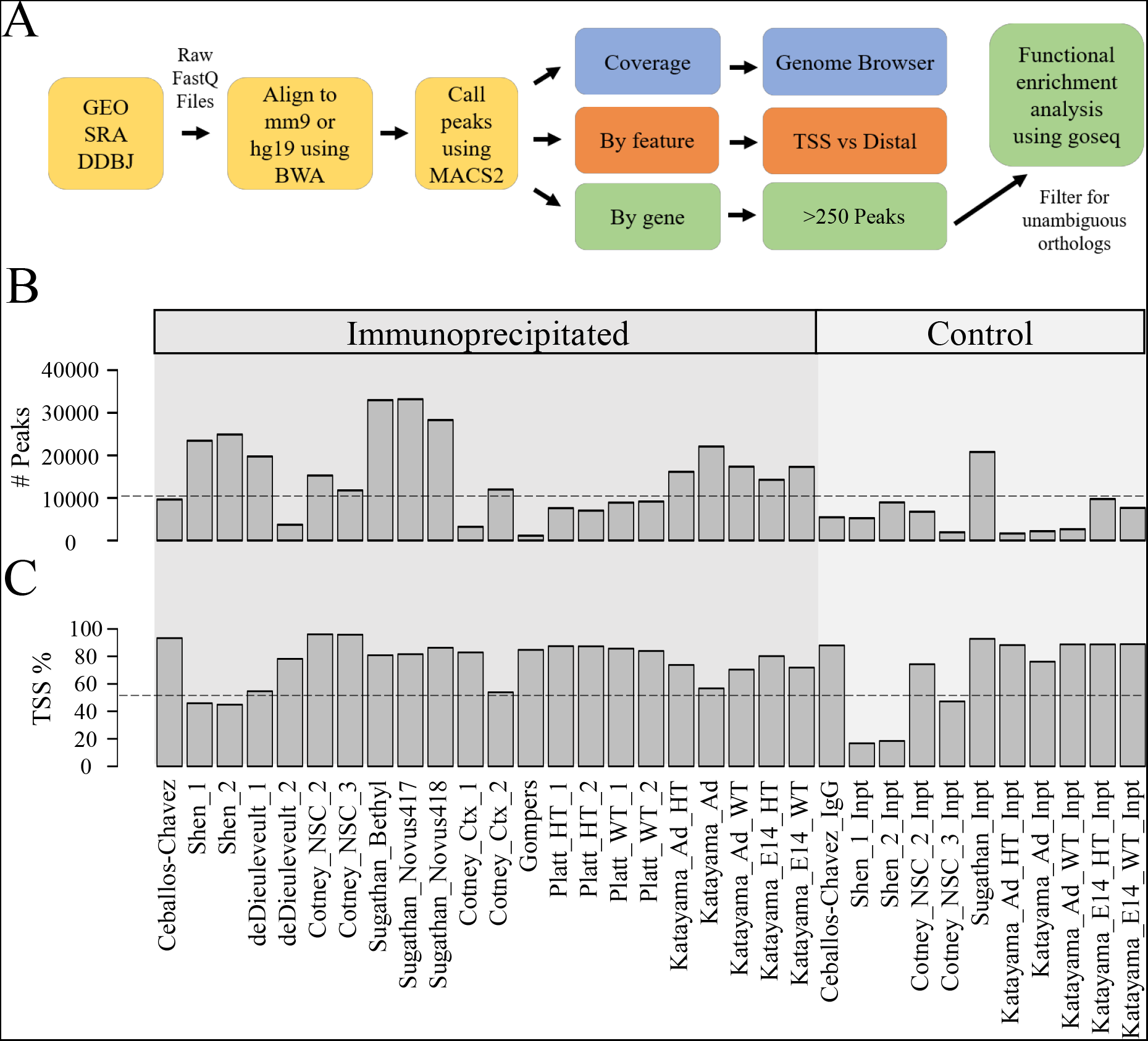
CHD8 binds to promoters across the genome. (A) ChIP-seq analysis pipeline. (B) Number of called peaks meeting a MACS2 significance of p < 0.00001. (C) Percentage of called peaks overlapping with the transcription start site of the nearest gene.

**Table 2.** Summary of datasets included in CHD8 ChIP-seq reanalysis. All datasets were performed using formaldehyde or another similar method of crosslinking before fragmentation and immunoprecipitation; E – embryonic day, P – postnatal day, h – human, NSCs-Neural Stem Cells, m – mouse, ESCs – Embryonic Stem Cells, HA -haemagglutinin, MNase – Micrococcal nuclease, *Chd8^+/-^ – CHD8* heterozygous mutation carrier.

| Fragmentation Method | Study | Model | Timepoint(s) | Tissue Collected | Antibody | Control |
| --- | --- | --- | --- | --- | --- | --- |
| Sonication | Cotney et al. 2015 | C57BL/6J mice; H9-derived hNSCs | E17.5; - | Frontal Cortex; - | αCHD8 (Abcam, ab114126) | Input |
| Sonication & Mnase | Katayama et al. 2016 | C57BL/6J mice (*Chd8*<sup>+/-</sup> & WT) | E14, P91 | Whole Brain | αCHD8 (Custom) | Input |
| Sonication | Platt et al. 2017 | C57BL/6J mice (*Chd8*<sup>+/-</sup> & WT) | P70-84 | Somatosensory Cortex | αCHD8 (Novus Biologicals, NB100-60417) | IgG |
| Sonication | Ceballos-Chavez et al. 2015 | hT47D-MTVL breast cancer cell | Before progestin stimulation | - | αCHD8 (Bethyl, A301-224A) | IgG |
| MNase | de Dieulevult et al. 2016 | mESCs with FLAG/HA-tagged CHD8 | - | - | αFLAG & αHA | Input |
| Sonication | Gompers et al. 2017 | C57BL/6N mice | ~P56 | Bulk Forebrain | αCHD8 (Abcam, ab114126) | Input |
| Sonication | Shen et al. 2015 | mRN2 cells | - | - | αCHD8 (Bethyl, A301-224A) | Input |
| Sonication | Sugathan et al. 2014 | iPSC-derived hNPCs | - | - | αCHD8 (Bethyl, A301-224A; Novus Biologicals, NB100-60417, NB100-60418) | Input |

Across all ChIP-seq datasets, CHD8 genomic interactions most commonly occurred near promoters (**Figure 3C**). Furthermore, binding to promoter-defined peaks tended to approach 100% as the number of called peaks decreased, suggesting that higher affinity interactions for CHD8 are largely at promoters. Increased affinity and frequency of promoter interactions by CHD8 was clearly evident in the coverage data signal for both mouse tissues (**Figure 4A**) and human cell lines (**Figure 4B**). For example, four genes encoding a transcription factor (*ADNP*), a chromatin remodeler (*SUV420H1*), a splicing factor (*TRA2B*), and a calcium binding protein important for cell cycle progression and ion channel signaling (*CALM2*) displayed CHD8 interactions almost exclusively near promoters.

**Figure 4.**
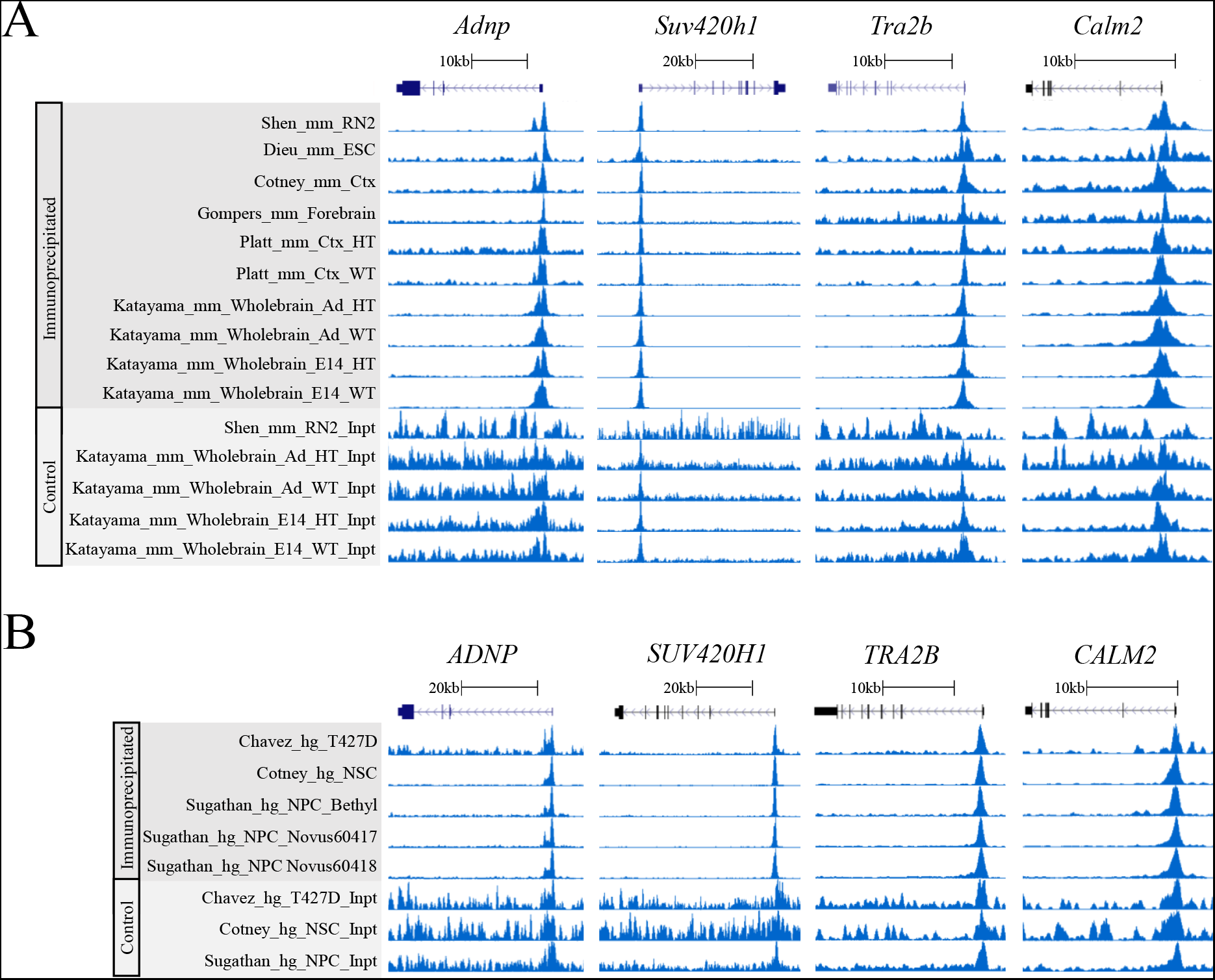
Examples of CHD8 binding near promoters of select chromatin (*ADNP, SUV420H1*), RNA processing (*TRA2B),* and neuronal function (*CALM2*) genes in the mouse (top) and human (**bottom**) ChIP-seq datasets.

*De novo* motif analysis performed on CHD8 peak regions across experiments identified various general promoter-associated transcription factor binding sequences, but no clear primary binding motif for CHD8 (**Supplementary Figure 4**). These findings are consistent with original publications, none of which identified a strong candidate primary binding motif, suggesting that CHD8 interactions are not mediated by direct DNA sequence recognition. Instead these results suggest that CHD8 genomic interaction specificity likely occurs through secondary interactions.

CHD8 GO analysis of the top 2000 called peaks ranked by signal strength and significance highlighted surprisingly consistent CHD8 interactions near homeostatic gene promoters at the FDR < 0.05 GO term association cutoff level (**Figure 5 top left**). When analyzing all called peaks, these terms were still enriched (**Figure 5 bottom left**). Analysis of all called peaks also identified CHD8 interactions with the expanded set of promoters that included genes associated with neuronal differentiation and function, as previously observed (**Figure 4**). However, unlike homeostatic gene targets, neuronal gene interaction was not statistically enriched across all relevant models suggesting the neuronal promoter interactions tended to have lower affinity and are not as a set enriched as CHD8 regulatory targets. Testing for the proportion of genes enriched for each GO term further indicated homeostatic gene promoter target specificity. We found significant enrichment of homeostatic promoter targets when analyzing the top 2000 peaks (**Figure 5 top right**). In contrast, almost all other GO terms were represented when considering all called peaks (**5 bottom right**). This suggested that while CHD8 has interactions throughout the genome with loci associated with various functions, homeostatic genes are consistent, high affinity CHD8 targets regardless of the model system.

**Figure 5.**
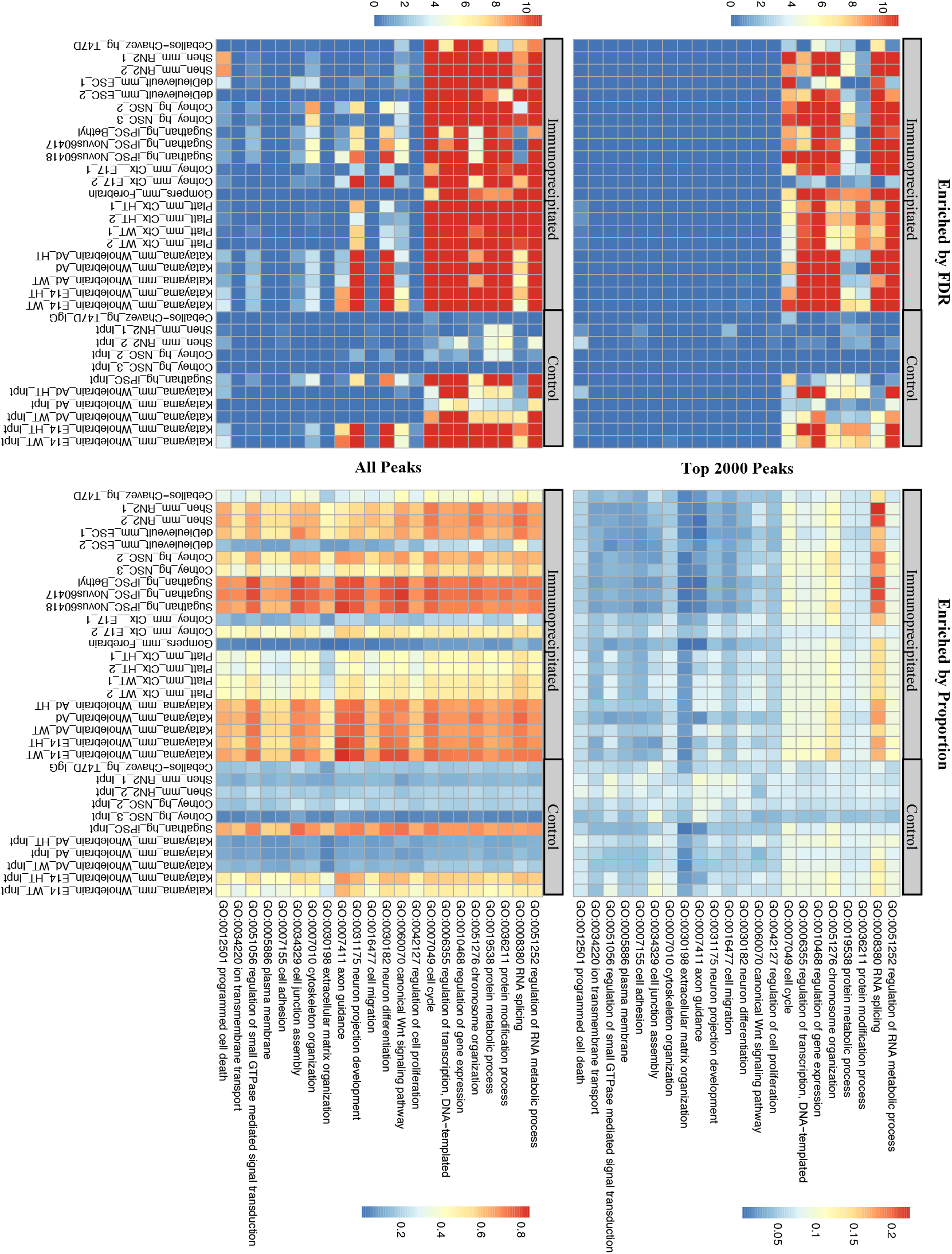
Unexplained specificity of CHD8 binding near gene regulator promoters. (**Left**) Enrichment of terms according to an FDR < 0.05 cutoff. (**Right**) Enrichment of terms according to proportion ranging from 0, not enriched, to 1, all genes in the term category. (**Top**) Analysis of the top 2000 significant peaks. (**Bottom**) Analysis of all peaks meeting a p < 0.00001 MACS2 significance level.

### Relationship between genomic interaction targets and gene expression changes across *CHD8* studies

Given that CHD8 targets a consistent set of promoters with a high level of affinity as well as an expanded set of loci with lower levels of affinity and *CHD8* knockdown or heterozygous knockout causes changes to gene expression, we tested whether CHD8 genomic interactions directly relate to changes in gene expression in *CHD8* models. Most genes with CHD8 interactions at or distal to the promoter did not exhibit significant changes in gene expression, regardless of the study, suggesting that there are additional determinants regarding sensitivity of regulatory target genes to *CHD8* dosage. While we did observe consistent patterns of overlap between CHD8 targets and downregulated DEGs, upregulated DEGs were not enriched across studies for CHD8 genomic interactions. Regardless of the experiment, CHD8 interaction affinity was also strongest for genes that were more highly expressed (**Supplementary Figure 5**).

For a subset of RNA-seq results, there was a strong overlap between CHD8 target genes from the ChIP-seq data and downregulated DEGs involved with cellular homeostasis. For example, when using the Gompers et al. (2017) RNA-seq data, there was an increased signature of downregulation in *Chd8* heterozygous mouse brain as CHD8 target affinity (i.e. ChlP-seq peak strength) increased (**Figure 6**). This was not unexpected considering homeostatic genes made up one of the two major signatures of differential gene expression we found and was the most strongly enriched set for CHD8 interaction. Eight out of the 18 analyzed datasets showed this overlap trend (**Supplementary Figure 6**). For instance, an increased signature of downregulation was also observed as CHD8 target affinity increased with *in vivo Chd8* knockdown in fetal mouse brain (Durak et al. 2016), though this RNA dataset had both neuronal and homeostatic terms enriched for DEGs (**Supplementary Figure 7**).

**Figure 6.**
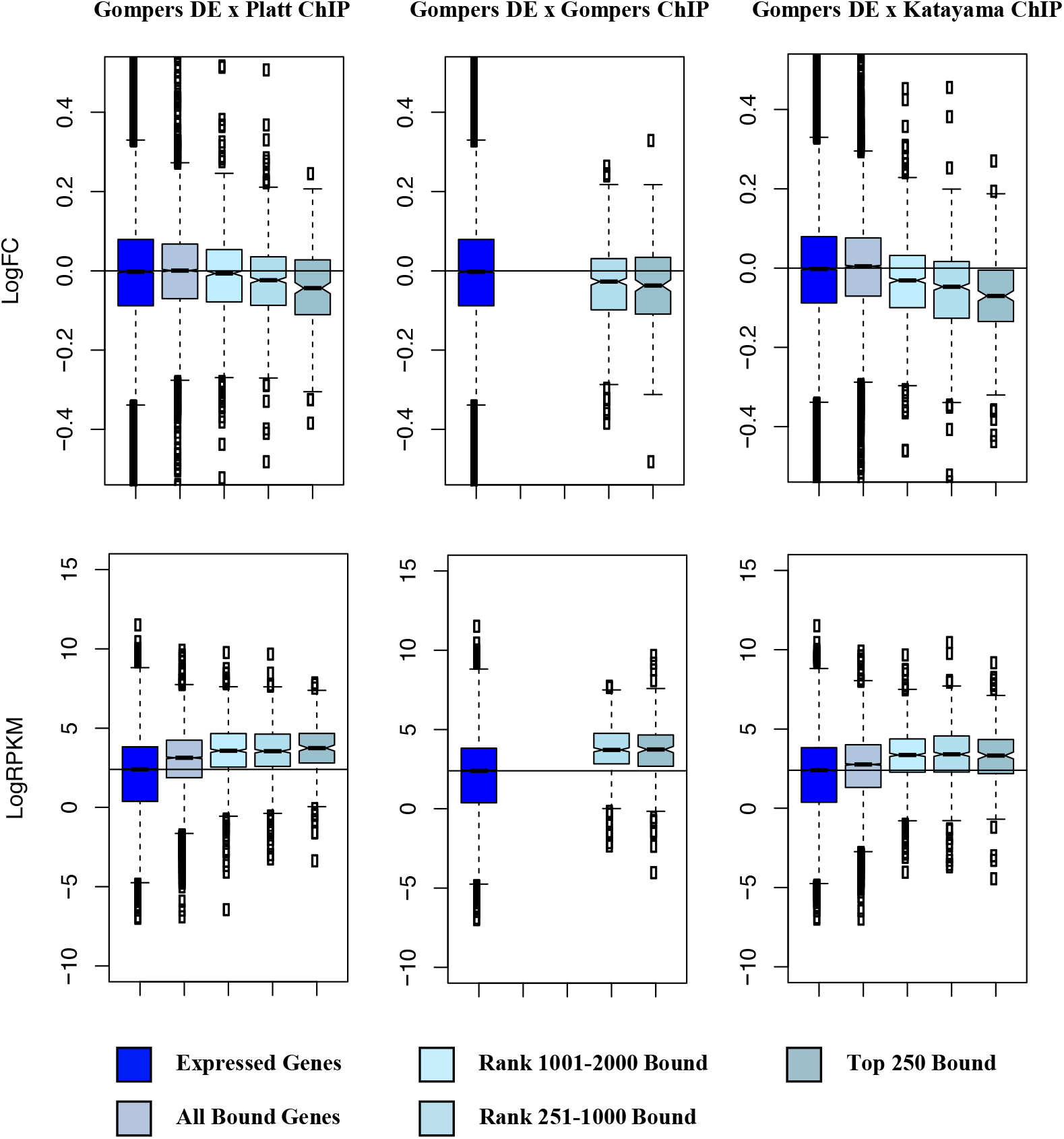
CHD8 regulates differentially expressed genes with high-confidence CHD8 binding. (**Left**) Comparison between the full Gompers et al. 2017 *Chd8* heterozygous mouse model differential expression gene set (DGE) and the Platt et al. 2017 Chd8 ChIP-seq dataset. (**Middle**) Comparison between the Gompers et al. DGE and Gompers et al. Chd8 ChIP-seq dataset. (**Right**) Comparison between the Gompers et al. DGE and Katayama et al. 2016 Chd8 ChIP-seq dataset. (**Top**) Change in expression of genes according to CHD8 binding compared to wild-type littermates. (**Bottom**) Change in sequence coverage of genes according to CHD8 binding. Boxes were plotted according to CHD8 binding affinity bins: all genes meeting a 1 count per million sequencing coverage threshold included in DEG analysis (Expressed Genes), any genes having CHD8 binding (All Bound Genes), and all genes having binding ranked according to CHD8 peak significance (Top 250 Bound, Rank 251-1000 Bound, Rank 1001-2000 Bound).

Genes associated with cellular homeostasis also tend to be at the high end of transcript expression level distributions, suggesting a relationship between highly expressed genes and dosage-sensitive CHD8 regulatory function. However, high levels of expression alone did not predict CHD8 interaction or DEG, indicating that expression level does not solely determine CHD8 interactions or sensitivity of regulatory targets to reduced *CHD8* dosage. This trend of negative correlation between CHD8 interaction affinity and changes in gene expression was less apparent with datasets having fewer than 500 differentially expressed genes and for the datasets (including many generated from *in vitro* models) where homeostasis gene expression signatures were not present (**Supplementary Figure 6, Figure 2**).

While our findings show model-specific variation, the patterns present across *CHD8* studies suggest a consistent relationship where reduced expression of *CHD8* leads to downregulation of CHD8 target genes associated with the cellular homeostasis signature, such as genes involved in cell cycle, chromatin organization, and RNA transcription and processing. These changes are seemingly stronger in *in vivo* models representing early stages of brain development, though they are still present in some models representing more mature brain tissue and cell types (**Figure 2**). In contrast, the observed differential expression of neuronal differentiation and neuronal function (e.g. synaptic) genes tends to occur in models representing more mature neuronal tissue or cell types and differentiated culture models inherently containing heterogenous cell populations at unknown stages of development.

## DISCUSSION

This meta-analysis of published genomic datasets from *in vitro* and *in vivo* mouse and human studies revealed both consistent and study-specific effects of *CHD8* haploinsufficiency on gene expression and largely concordant high-affinity CHD8 genomic interaction loci. Knockdown or heterozygous mutation of *CHD8* led to characteristic changes in gene expression across studies and model systems. At the gene-by-gene level, these expression changes varied considerably between *CHD8* models. However, at the level of gene set enrichment, we found global patterns of transcriptional dysregulation of genes involved in cellular homeostasis and neuronal development and function. Comparison across ChIP-seq experiments shows that CHD8 preferentially targets promoters, with no evidence of direct binding through a specific DNA motif. Surprisingly, we found that peaks with the highest signal were constant across experiments, regardless of the model, suggesting that CHD8 preferentially interacts with promoters of a set of genes linked to processes involved in cellular homeostasis, genome function, and RNA processing. Our findings strongly support signatures of reduced transcription of CHD8 target genes in the studied models that were dependent on dosage, though our data also highlight widespread genomic promoter interactions for CHD8 without obvious strong impacts to most targets. We verified the presence of changes to gene expression specific to neuronal differentiation and function following *CHD8* haploinsufficiency across studies, however, these changes do not appear to be through direct disruption to neural cell-type or stage-specific CHD8 regulatory activity via high-affinity interactions with relevant promoters. While the clear concordance in high-affinity genomic CHD8 interactions suggests common regulatory functions across cell types, it remains to be examined whether the observed dysregulation of neuronal genes is related to context-dependent CHD8 regulatory activity in the brain given the current cellular heterogeneity and technical challenges existing with available CHD8 ChIP-seq. Our results illustrate the power and limitations of comparing genomic datasets and challenge previous assumptions regarding the regulatory mechanisms and transcriptional pathology associated with *CHD8* haploinsufficiency.

We note a number of technical issues that impacted this meta-analysis, many of which are associated with variation in methods and sequencing depth. Surprisingly, we found considerable differences in *CHD8* expression across models despite the common design of testing the impacts of haploinsufficiency. Though we did not find an obvious correlation between *CHD8* transcript levels and up- or downregulated gene expression, it seems likely that differences in experimental design, including *CHD8* knockdown or knockout, contributed toward meaningful variation between models. Changes to *CHD8* dosage have been shown to have strong and potentially opposing effects on cellular function. For instance, homozygous knockout of *CHD8* has been described to cause severe developmental arrest and widespread apoptosis leading to early embryonic lethality (Nishiyama et al. 2009) while heterozygous mutation can lead to increases in proliferation (Gompers et al. 2017). These are important considerations for interpreting studies of haploinsufficiency, as allelic or genetic background effects as well as variation in transcriptional knockdown with shRNA constructs may have significant biological consequences. At a minimum, consistent measures of *CHD8* knockdown or haploinsufficiency, such as measuring transcript-level mRNA levels with RNA-seq or protein levels with a standardized antibody and methodology, should be a goal for future studies to enable comparison across publications. We also noted differences in ChIP-seq datasets that hindered comparisons. For example, enrichment in control libraries was present across several published datasets. Different studies also used various CHD8 antibodies with unknown and unvalidated CHD8 specificities. Nonetheless, by examining patterns across datasets, we identified consistent patterns of enrichment suggesting that overall findings from ChIP-seq targeting CHD8 could reliably identify high affinity interactions.

Despite the limitations of comparing genomic datasets across variable models, our analysis challenges two simple models regarding pathological mechanisms of *CHD8* haploinsufficiency. The first model the transcriptional signatures present across studies refute is that pathology due to *CHD8* haploinsufficiency is primarily due to alterations in patterning during early brain development. While our meta-analysis clearly supports impacts to proliferation and neuronal differentiation consistent with published findings on proliferation and brain volume (Bernier et al. 2014, Gompers et al. 2017, Katayama et al. 2016, Platt et al. 2017, Durak et al. 2016), we also observed evidence of dysregulation of genes involved in mature neuron function, including synaptic genes. This is consistent with observation that CHD8 is still highly expressed in adulthood (Gompers et al. 2017, Platt et al. 2017, Maussion et al. 2015), that mutations to *CHD8* continue to lead to differential gene expression and behavioral phenotypes in adult mice (Gompers et al. 2017, Katayama et al. 2016, Platt et al. 2017), and with limited evidence of synaptic dysfunction associated with *Chd8* haploinsufficiency (Platt et al. 2016). Further work will be required to establish the role and requirement for CHD8 in mature neurons and other cell types in the brain.

Second, the signatures present in this meta-analysis suggest that pathology observed in *CHD8* models and patients with *CHD8* mutations is not due to targeted impacts to specific populations of cell-types or due to impacts limited to specific brain regions. In this analysis of many individual datasets, CHD8 had genomic interactions near promoters of genes important for cellular homeostasis and neuronal development and function that were enriched in the transcriptomic analysis. However, only homeostasis genes were characterized as high affinity CHD8 targets and tended to be sensitive to decreases in *CHD8* expression and haploinsufficiency. Despite evidence that these genes are not high-affinity CHD8 targets, we did observe enrichment of differentially expressed neuronal genes in the CHD8 interaction analysis. One explanation for this finding is that *CHD8* haploinsufficiency indirectly causes large-scale dysregulation of neuronal genes via disruptions to upstream transcriptional regulators that are direct CHD8 targets. This would explain the appearance of cell-type-specific transcriptional changes in the absence of actual cell-type specific CHD8 function. Nonetheless, given the technical limitations of current studies we cannot rule out the possibility of cell-type or context-dependent specificity of CHD8 function.

It is clear from previous publications and this meta-analysis that CHD8 is critical for neurodevelopment. Our results suggest that CHD8 functions to regulate cellular homeostasis required for genomic control of proliferation and differentiation. As an essential gene with widespread expression across neuronal and glial cell types, homozygous loss of *CHD8* may impact cellular viability in general, while heterozygous mutation or knockdown might have subtler, context-specific impacts. Such a model would explain the widespread changes in gene expression across model systems and varied reports of impact on proliferation depending on dosage. Our results raise two questions that could be addressed by application of RNA-seq and ChIP-seq in the future: 1) What are the developmental stage, cell-type, and region-specific impacts of *CHD8* haploinsufficiency in the developing and mature brain, and 2) Does CHD8 have context-dependent function in specific stages, cell types, and regions with regard to genomic interaction patterns? Beyond addressing these two key issues, additional clarity regarding the role of CHD8 in the brain will come from studies examining the molecular interaction partners and impacts on chromatin, transcription, and RNA processing. As *CHD8* haploinsufficiency may represent common features of haploinsufficiency of other general chromatin remodelers implicated in patient studies, further characterization of *CHD8* models and CHD8 genomic interactions could reveal essential functions driving pathology in neurodevelopmental disorders.

## AUTHOR CONTRIBUTIONS

AW and AN conceived of the project. AW, KL, and AN performed analysis of RNA-seq experiments. AW, RCP, and AN performed analysis of ChIP-seq experiments. AW and AN drafted the manuscript. All authors contributed to manuscript revision.

## ACKNOWLEDGEMENTS

The authors would like to thank the authors from the original *CHD8* studies who provided access to the raw data for analysis.

---

**Supplementary Figure 1.**
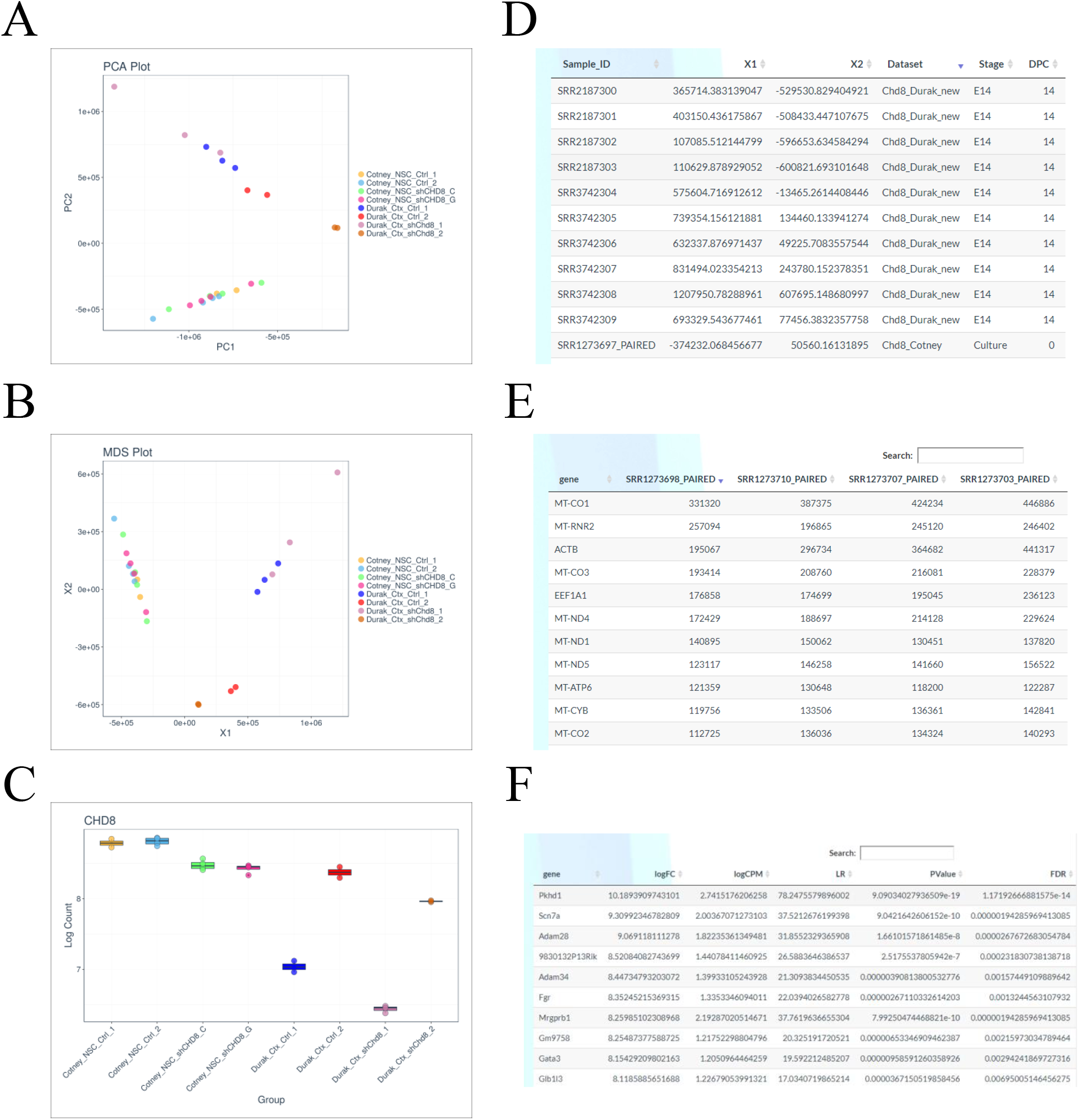
Analyzing individual *CHD8* model gene expression and pairwise comparisons through the Shiny interactive web browser. (**A**) The *CHD8* RNA-seq Shiny app can generate a principle component analysis (PCA) scatter plot with any of the RNA-seq datasets loaded onto the app. You can tailor the plot according to several parameters including sample number, principle component, gene of interest, dataset, timepoint, sex, model organism, mouse or cell line, and genotype. This plot shows the Cotney et al. 2015 and Durak et al. 2016 RNA-seq datasets plotted according to the experimental design, which is in this case the control and experimental constructs. A multidimensional scaling (MDS) plot (**B**) and log fold change differential gene expression bar plot (**C**) generated using Shiny is also shown with experimental design chosen as the display parameters. Almost all of the same display parameters for the PCA plot are available for the MDS and box plots. Supplementary tables showing metadata (**D**) and gene counts (**E**) are also available through the Shiny app. (**F**) Table showing log fold gene expression changes and significance values for individual genes between the Cotney et al. 2015 and Durak et al. 2016 RNA-seq datasets. Heat maps and scatter plots of gene expression changes are also available. All plots and tables generated using Shiny can be downloaded from the app. Datasets can be analyzed using pseudo counts or relative expression.

Supplementary Figure 2

(See Supplementary Figure File)

Full list of terms from the gene ontology analysis using goseq. Terms were selected from this list to create Figure 2. All RNA-seq datasets were included in this figure. All terms included met an FDR < 0.05 cutoff.

**Supplementary Figure 3.**
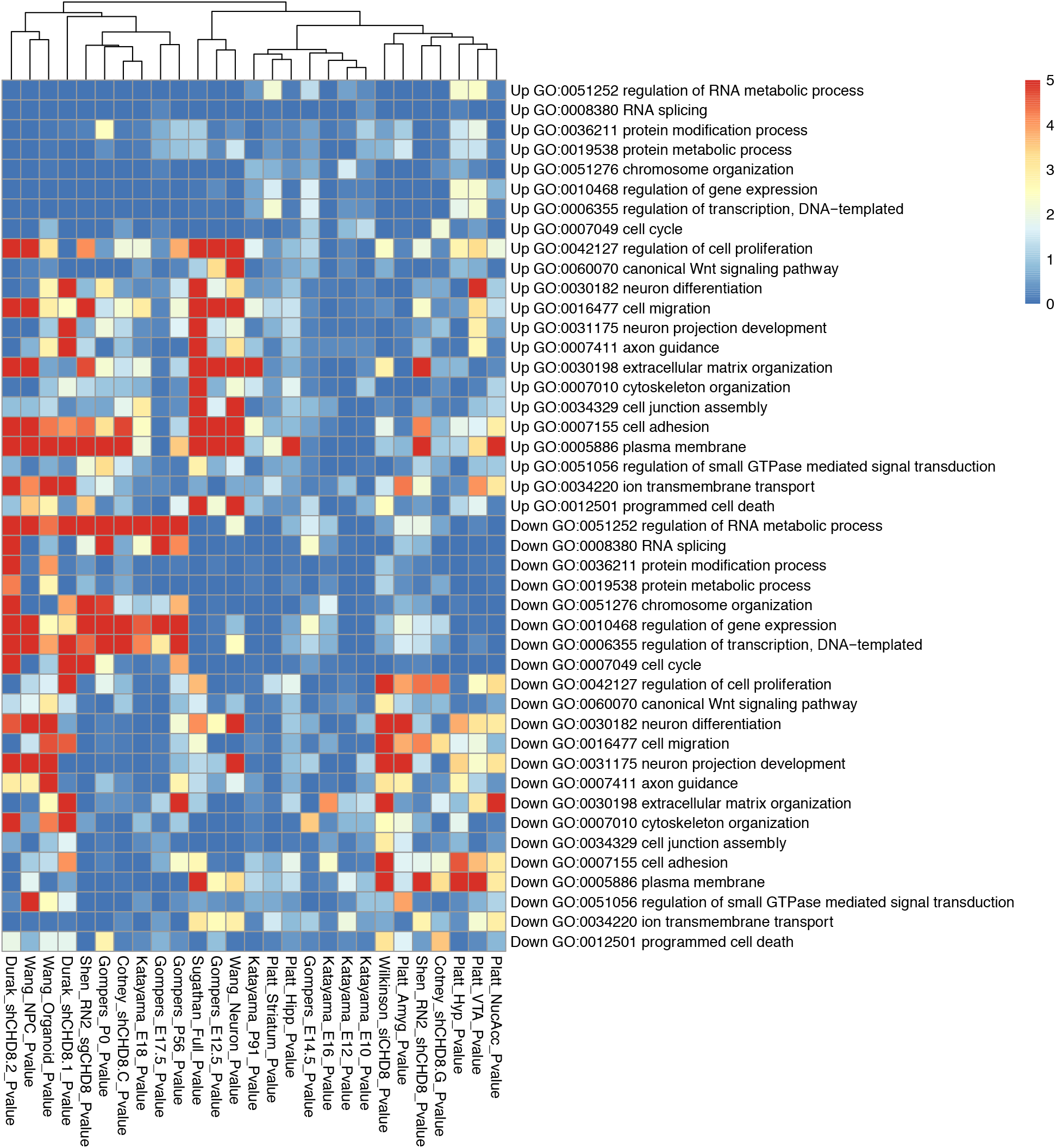
Enrichment of gene regulation and neurodevelopmental ontology terms meeting a goseq p < 0.05 significance level in all Up- and Down-regulated datasets. (**Top**) Upregulated gene ontology enrichment. (**Bottom**) Downregulated gene ontology enrichment.

**Supplementary Figure 4.**
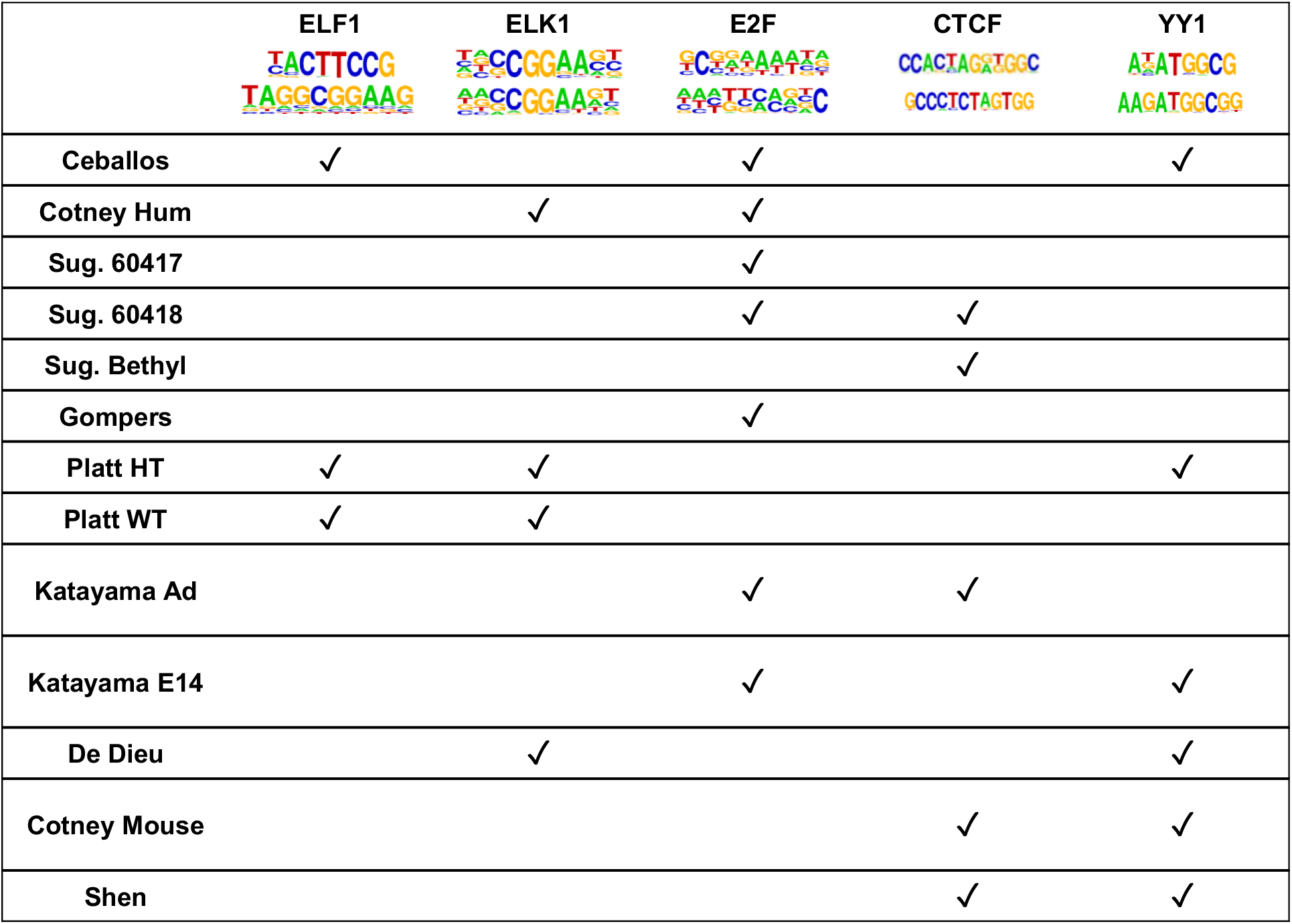
No obvious primary motif associated with CHD8 binding. Each dataset was analyzed using HOMER to look for common motifs enriched in CHD8 ChIP-seq datasets. ELF1, ELK1, E2F, CTCF, and YY1 transcription factors were the motifs that were commonly represented across datasets. Ceballos - Ceballos-Chavez et al. 2015 ChIP-seq dataset, Sug. 60417, 60418, Bethyl - Sugathan et al. 2014 ChIP-seq datasets split according to antibody used (60417, NB100-60417; 60418, NB100-60418), Gompers - Gompers et al. 2017 ChIP-seq dataset, Platt HT, WT - Platt et al. 2017 ChIP-seq datasets split according to genotype of the samples (HT, heterozygous; WT, wild-type), Katayama - Katayama et al. 2016 ChIP-seq datasets split according to age of the samples (Ad, postnatal day 91; E14, embryonic day 14), De Dieu - De Dieulevuelt et al. 2016 ChIP-seq dataset, Cotney Hum, Mouse - Cotney et al. 2015 ChIP-seq datasets split according to model organism (Hum, human), Shen - Shen et al. 2015 ChIP-seq dataset.

**Supplementary Figure 5.**
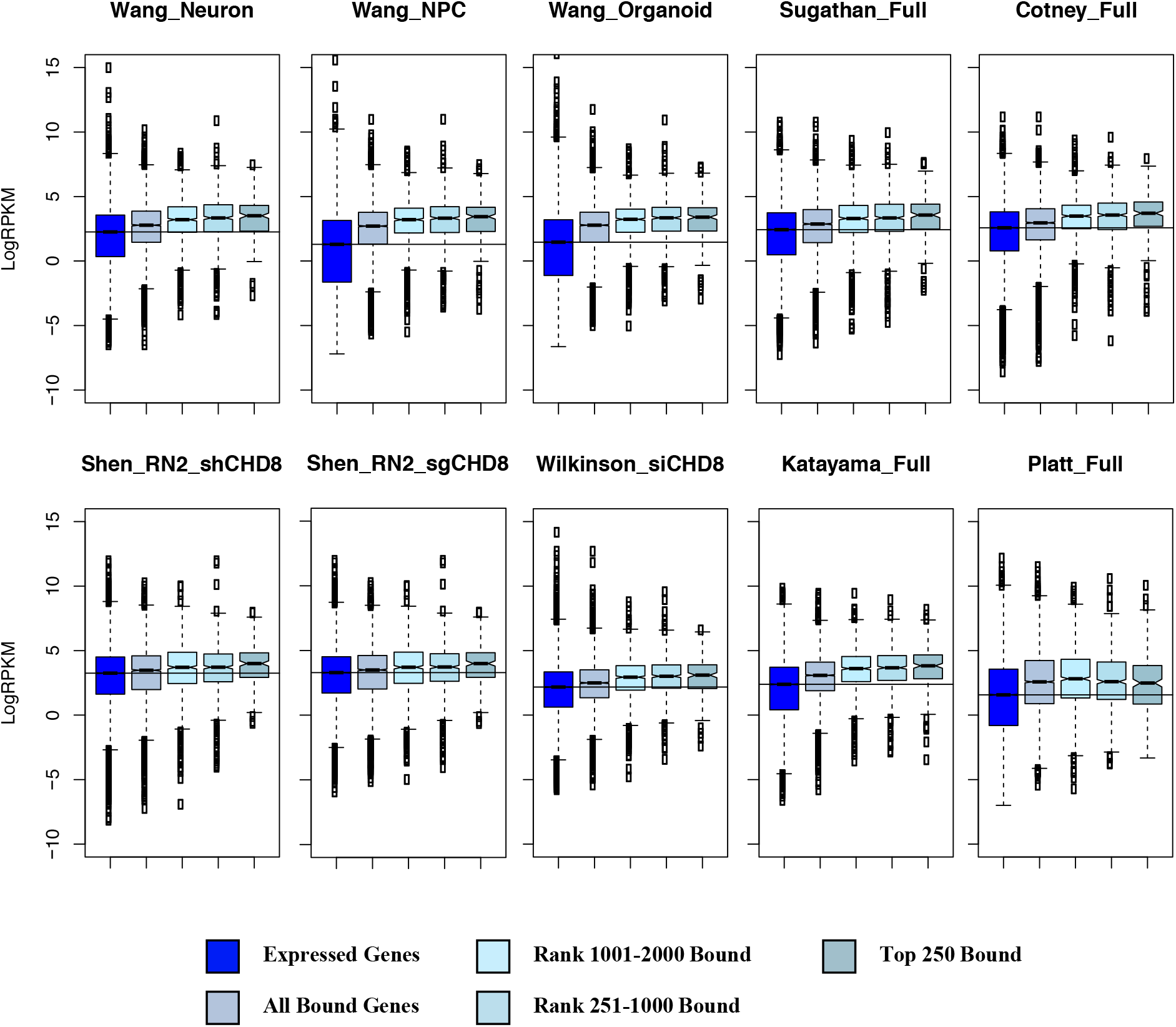
Chd8 regulates highly expressed genes. Each dataset is labelled showing changes in sequencing coverage according to changes in CHD8 binding affinity. The Chd8 ChIP-seq dataset used was from Platt et al. 2017. Full models for each dataset were chosen as they exhibited similar signal as the individual timepoint or brain region datasets.

**Supplementary Figure 6.**
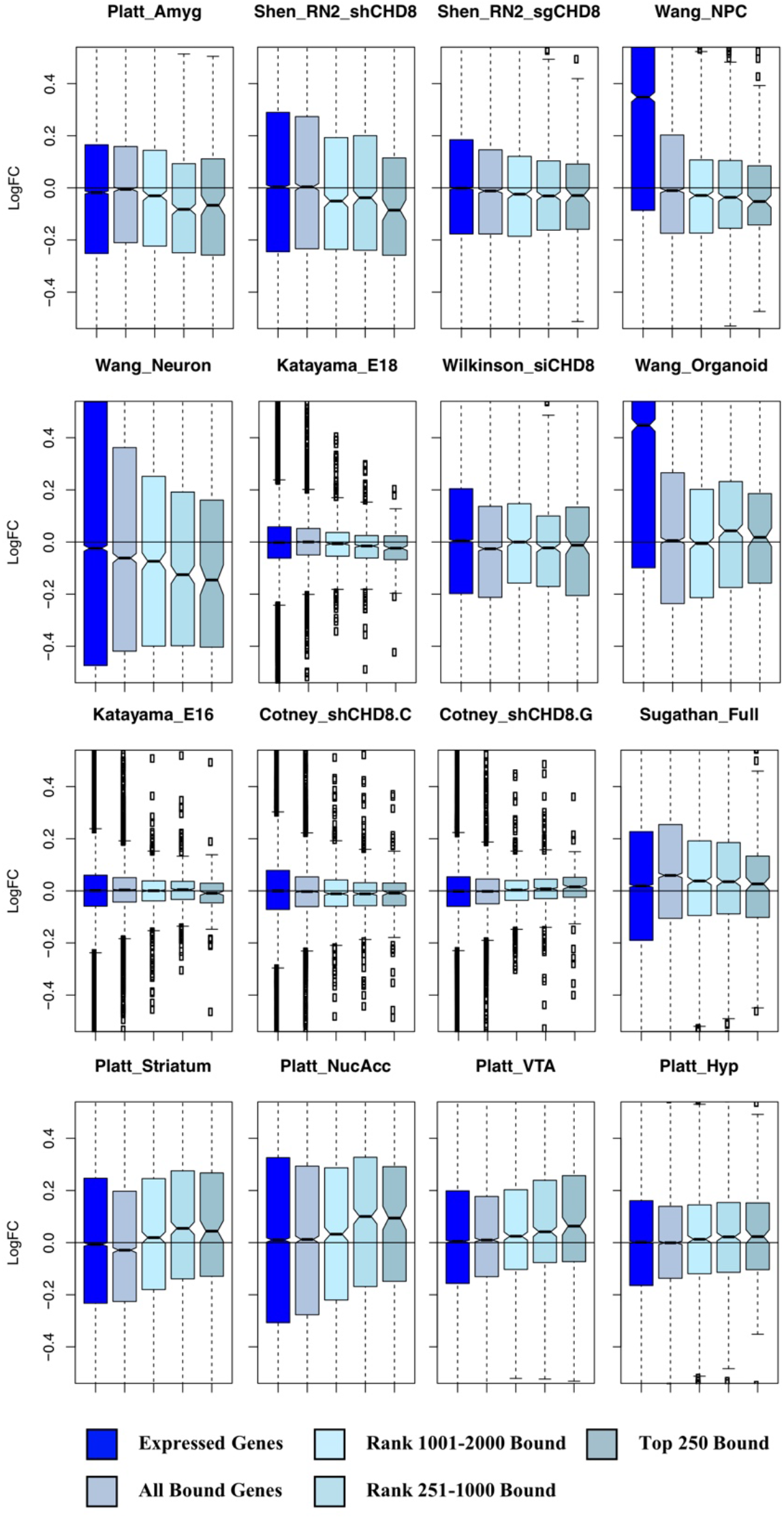
Remaining fold change plots from the CHD8 binding by differential gene expression comparison analysis. All datasets were analyzed using the Platt et al. 2017 Chd8 ChIP-seq dataset. Datasets are loosely organized based on overlap between downregulated genes, no clear trend, or upregulated genes from top to bottom, which sometimes spanned multiple rows.

**Supplementary Figure 7.**
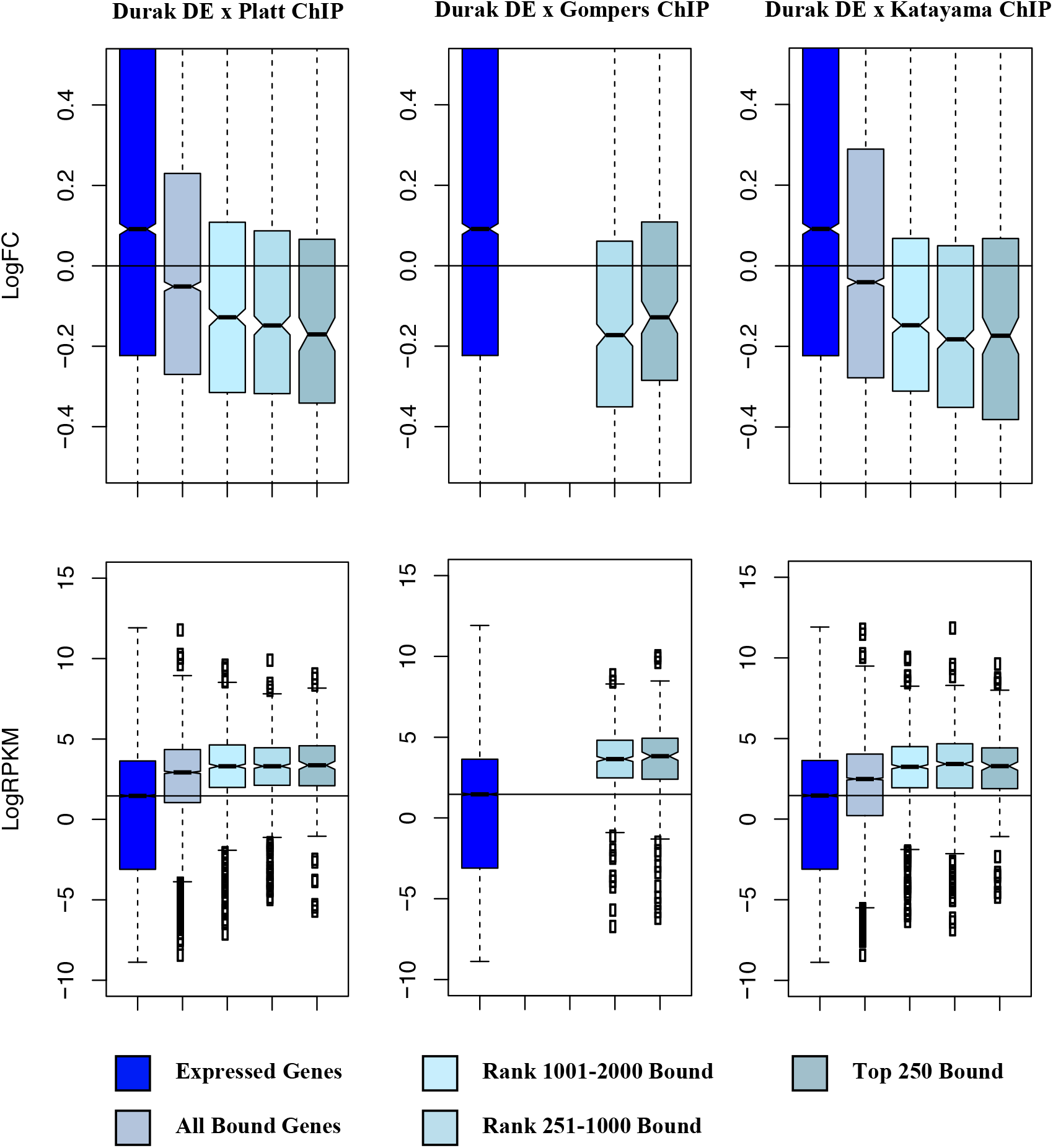
Chd8 regulates genes differentially expressed with *Chd8* knockdown. (**Left**) Comparison between the full Durak et al. 2016 *Chd8* knockdown mouse model differential expression gene set (DGE) and the Platt et al. 2017 Chd8 ChIP-seq dataset. (**Middle**) Comparison between the Durak et al. DGE and Gompers et al. Chd8 ChIP-seq dataset. (**Right**) Comparison between the Durak et al. DGE and Katayama et al. 2016 Chd8 ChIP-seq dataset. (**Top**) Change in expression of all genes compared to wild-type littermates according to changes in CHD8 binding affinity. (**Bottom**) Changes in sequencing coverage of genes according to changes in CHD8 binding affinity.

## References

Barnard, R. A., Pomaville, M. B., and O’Roak, B. J. (2015). Mutations and modeling of the chromatin remodeler CHD8 define an emerging autism etiology. Front. Neurosci. 9, 477. doi: 10.3389/fnins.2015.00477

Bernier, R., Golzio, C., Xiong, B., Stessman, H., Coe, B., Penn, et al. (2014). Disruptive CHD8 mutations define a subtype of autism early in development. Cell 158, 263–276. doi: 10.1016/j.cell.2014.06.017

Ceballos-Chávez, M., Subtil-Rodríguez, A., Giannopoulou, E., Soronellas, D., Vázquez-Chávez, E., Vicent, G., et al. (2015). The chromatin remodeler CHD8 is required for activation of progesterone receptor-dependent enhancers. PLoS Genet. 11, e1005174. doi: 10.1371/journal.pgen.1005174

Cotney, J., Muhle, R., Sanders, S., Liu, L., Willsey, A., Niu, W., et al. (2015). The autism-associated chromatin modifier CHD8 regulates other autism risk genes during human neurodevelopment. Nat. Comm. 6, 6404. doi: 10.1038/ncomms7404

de Dieuleveult, M., Yen, K., Hmitou, I., Depaux, A., Boussouar, F., Dargham, D., et al. (2016). Genome-wide nucleosome specificity and function of chromatin remodellers in ES cells. Nature 530, 113–116. doi: 10.1038/nature16505

De Rubeis, S., He, X., Goldberg, A., Poultney, C., Samocha, K., Ercument Cicek, A., et al. (2014). Synaptic, transcriptional and chromatin genes disrupted in autism. Nature 515, 209–215. doi: 10.1038/nature13772

Dobin, A., Davis, C. A., Schlesinger, F., Drenkow, J., Zaleski, C., Jha, S., et al. (2013). STAR: ultrafast universal RNA-seq aligner. Bioinformatics 29, 15–21. doi: 10.1093/bioinformatics/bts635

Durak, O., Gao, F., Kaeser-Woo, Y., Rueda, R., Martorell, A., Nott, A., et al. (2016). Chd8 mediates cortical neurogenesis via transcriptional regulation of cell cycle and Wnt signaling. Nat. Neurosci. 19, 1477–1488. doi: 10.1038/nn.4400

Fang, M., Hutchinson, L., Deng, A., and Green, M. R. (2016). Common BRAF(V600E)-directed pathway mediates widespread epigenetic silencing in colorectal cancer and melanoma. PNAS 113, 1250–1255. doi: 10.1073/pnas.1525619113

Feng, J., Liu, T., and Zhang, Y. (2011). Using MACS to identify peaks from ChIP-Seq data. Curr. Protoc. Bioinformatics Chapter 2, Unit 2.14. doi: 10.1002/0471250953.bi0214s34

Flanagan, J., Mi, L., Chruszcz, M., Cymborowski, M., Clines, K., Kim, Y., et al. (2005). Double chromodomains cooperate to recognize the methylated histone H3 tail. Nature 438, 1181–1185. doi: 10.1038/nature04290

Gompers, A.L., Su-Feher, L., Ellegood, J., Copping, N.A., Riyadh, A., Stradleigh, T.W., et al. (2017). Germline Chd8 haploinsufficiency alters brain development in mouse. Nat. Neurosci. 20, 1062–1073. doi: 10.1038/nn.4592

Hall, J., and Georgel, P. (2007). CHD proteins: a diverse family with strong ties. Biochem. Cell Biol. 85, 463–476. doi: 10.1139/O07-063

Han, H., Braunschweig, U., Gonatopoulos-Purnatzis, T., Weatheritt, R.J., Hirsch, C.L., Ha, K.C., et al. (2017). Multilayered Control of Alternative Splicing Regulatory Networks by Transcription Factors. Mol. Cell 65, 539–553. doi: 10.1016/j.molcel.2017.01.011

Hargreaves, D., and Crabtree, G. (2011). ATP-dependent chromatin remodeling: genetics, genomics and mechanisms. Cell Res. 21, 396–420. doi: 10.1038/cr.2011.32

Heinz, S., Benner, C., Spann, N., Bertolino, E., Lin, Y. C., Laslo, P., et al. (2010). Simple combinations of lineage-determining transcription factors prime cis-regulatory elements required for macrophage and B cell identities. Mol. Cell 38, 576–589. doi: 10.1016/j.molcel.2010.05.004

Iossifov, I., O’Roak, B. J., Sanders, S. J., Ronemus, M., Krumm, N., Levy, D., et al. (2014). The contribution of de novo coding mutations to autism spectrum disorder. Nature 515, 216–221. doi: 10.1038/nature13908

Ishihara, K., Oshimura, M., and Nakao, M. (2006). CTCF-dependent chromatin insulator is linked to epigenetic remodeling. Mol. Cell 23, 733–742. doi: 10.1016/j.molcel.2006.08.008

Katayama, Y., Nishiyama, M., Shoji, H., Ohkawa, Y., Kawamura, A., Sato, T., et al. (2016). CHD8 haploinsufficiency results in autistic-like phenotypes in mice. Nature 537, 675–679. doi: 10.1038/nature19357

Krumm, N., O’Roak, B., Shendure, J., and Eichler, E. (2014). A de novo convergence of autism genetics and molecular neuroscience. Trend Neurosci. 37, 95–105. doi: 10.1016/j.tins.2013.11.005

Li, H., and Durbin, R. (2009). Fast and accurate short read alignment with Burrows-Wheeler transform. Bioinformatics 25, 1754–1760. doi: 10.1093/bioinformatics/btp324

Liao, Y., Smyth, G. K., and Shi, W. (2014). featureCounts: an efficient general purpose program for assigning sequence reads to genomic features. Bioinformatics 30, 923–930. doi: 10.1093/bioinformatics/btt656

Marfella, C., and Imbalzano, A. (2007). The Chd family of chromatin remodelers. Mutat. Res. Fund. Mol. Mech. Mut. 618, 30–40. doi: 10.1016/j.mrfmmm.2006.07.012

Marinov, G. K., Kundaje, A., Park, P. J., and Wold, B. J. (2014). Large-scale quality analysis of published ChIP-seq data. G3 4, 209–223. doi: 10.1534/g3.113.008680

Maussion, G., Diallo, A.B., Gigek, C. O., Chen, E. S., Crapper, L., Théroux, J. F., et al. (2015). Investigation of genes important in neurodevelopment disorders in adult human brain. Hum. Genet. 134, 1037–1053. doi: 10.1007/s00439-015-1584-z

McCarthy, S. E., Gillis, J., Kramer, M., Lihm, J., Yoon, S., Berstein, Y., et al. (2014). De novo mutations in schizophrenia implicate chromatin remodeling and support a genetic overlap with autism and intellectual disability. Mol. Psych. 19, 652–658. doi: 10.1038/mp.2014.29

McKnight, J., Jenkins, K., Nodelman, I., Escobar, T., and Bowman, G. (2011). Extranucleosomal DNA binding directs nucleosome sliding by Chd1. Mol. Cell Biol. 31, 4746–4759. doi: 10.1128/MCB.05735-11

Nishiyama, M., Oshikawa, K., Tsukada, Y., Nakagawa, T., Iemura, S., Natsume, T., et al. (2009). CHD8 suppresses p53-mediated apoptosis through histone H1 recruitment during early embryogenesis. Nat. Cell Biol. 11, 172–182. doi: 10.1038/ncb1831

O’Roak, B., Vives, L., Fu, W., Egertson, J., Stanaway, I., Phelps, I., et al. (2012a). Multiplex targeted sequencing identifies recurrently mutated genes in autism spectrum disorders. Science 338, 1619–1622. doi: 10.1126/science.1227764

O’Roak, B., Vives, L., Girirajan, S., Karakoc, E., Krumm, N., Coe, B., et al. (2012b). Sporadic autism exomes reveal a highly interconnected protein network of de novo mutations. Nature 485, 246–250. doi: 10.1038/nature10989

Parikshak, N., Luo, R., Zhang, A., Won, H., Lowe, J., Chandran, V., et al. (2013). Integrative functional genomic analyses implicate specific molecular pathways and circuits in autism. Cell 155, 1008–1021. doi: 10.1016/j.cell.2013.10.031

Platt, R., Zhou, Y., Slaymaker, I., Shetty, A., Weisbach, N., Kim, J., et al. (2017). Chd8 mutation leads to autistic-like behaviors and impaired striatal circuits. Cell Rep. 19, 335–350. doi: 10.1016/j.celrep.2017.03.052

Robinson, M. D., McCarthy, D. J., and Smyth, G. K. (2010). edgeR: a Bioconductor package for differential expression analysis of digital gene expression data. Bioinformatics 26, 139–140. doi: 10.1093/bioinformatics/btp616

Rodriguez-Paredes, M., Ceballos-Chavez, M., Esteller, M., Garcia-Dominguez, M., and Reyes, J. (2009). The chromatin remodeling factor CHD8 interacts with elongating RNA polymerase II and controls expression of the cyclin E2 gene. Nucl. Acid Res. 37, 2449–2460. doi: 10.1093/nar/gkp101

Sanders, S. J., He, X., Willsey, A. J., Ercan-Sencicek, A. G., Samocha, K. E., Cicek, A. E., et al. (2015). Insights into autism spectrum disorder genomic architecture and biology from 71 risk loci. Neuron 87, 1215–1233. doi: 10.1016/j.neuron.2015.09.016

Shen, C., Ipsaro, J. J., Shi, J., Milazzo, J. A., Wang, E., Roe, J.-S., et al. (2015). NSD3-short is an adaptor protein that couples BRD4 to the CHD8 chromatin remodeler. Mol. Cell 60, 847–859. doi: 10.1016/j.molcel.2015.10.033

Sugathan, A., Biagioli, M., Golzio, C., Erdin, S., Blumenthal, I., Manavalan, P., et al. (2014). CHD8 regulates neurodevelopmental pathways associated with autism spectrum disorder in neural progenitors. P.N.A.S. 111, E4468–E4477. doi: 10.1073/pnas.1405266111

Tatton-Brown, K., Loveday, C., Yost, S., Clarke, M., Ramsay, E., Zachariou, A., et al. (2017). Mutations in epigenetic regulation genes are a major cause of overgrowth with intellectual disability. Am. J. Hum. Genet. 100, 725–736. doi: 10.1016/j.ajhg.2017.03.010

Thompson, B., Tremblay, V., Lin, G., and Bochar, D. (2008). CHD8 is an ATP-Dependent chromatin remodeling factor that regulates beta-catenin target genes. Mol. Cell Biol. 28, 3894–3904. doi: 10.1128/MCB.00322-08

Tong, J.K., Hassig, C.A., Schnitzler, G.R., Kingston, R.E., and Schreiber, S.L. (1998). Chromatin deacetylation by an ATP-dependent nucleosome remodeling complex. Nature 395, 917–921. doi: 10.1038/27699

Vissers, L. E. L. M., Gilissen, C., and Veltman, J. A. (2016). Genetic studies in intellectual disability and related disorders. Nat. Rev. Genet. 17, 9–18. doi: 10.1038/nrg3999

Wang, L., Wang, S., and Li, W. (2012). RSeQC: quality control of RNA-seq experiments. Bioinformatics 28, 2184–2185. doi: 10.1093/bioinformatics/bts356

Wang, P., Lin, M., Pedrosa, E., Hrabovsky, A., Zhang, Z., Guo, W., et al. (2015). CRISPR/Cas9-mediated heterozygous knockout of the autism gene CHD8 and characterization of its transcriptional networks in neurodevelopment. Mol. Autism 6, 55. doi: 10.1186/s13229-015-0048-6

Wang, P., Mokhtari, R., Pedrosa, E., Kirschenbaum, M., Bayrak, C., Zheng, D., et al. (2017). CRISPR/Cas9-mediated heterozygous knockout of the autism gene CHD8 and characterization of its transcriptional networks in cerebral organoids derived from iPS cells. Mol. Autism 8, 11. doi: 10.1186/s13229-017-0124-1

Wilkinson, B., Grepo, N., Thompson, B. L., Kim, J., Wang, K., Evgrafov, O. V., et al. (2015). The autism-associated gene chromodomain helicase DNA-binding protein 8 (CHD8) regulates noncoding RNAs and autism-related genes. Transl. Psych. 5, e568. doi: 10.1038/tp.2015.62

Young, M. D., Wakefield, M. J., Smyth, G. K., and Oshlack, A. (2010). Gene ontology analysis for RNA-seq: accounting for selection bias. Genome Biol. 11, R14. doi: 10.1186/gb-2010-11-2-r14

Yuan, C., Zhao, X., Florens, L., Swanson, S., Washburn, M., and Hernandez, N. (2007). CHD8 associates with human Staf and contributes to efficient U6 RNA polymerase III transcription. Mol. Cell Biol. 27, 8729–8738. doi: 10.1128/MCB.00846-07

